# Ecological associations of the coastal marsh periwinkle snail *Littoraria irrorata*: field and laboratory evidence of vegetation habitat preferences

**DOI:** 10.1101/2024.11.24.625093

**Authors:** David H. Klinges, Charles W. Martin, Brian J. Roberts

## Abstract

Coastal salt marshes serve as the margin between terrestrial and marine biomes, provide a variety of important services, and are dynamic ecosystems characterized by keystone species that shape trophic networks. In coastal salt marshes of the Eastern Atlantic and Gulf of Mexico, marsh periwinkle snails (*Littoraria irrorata*) exhibit high abundance and form critical trophic pathways as important herbivores and detritivores. Specifically, snails forage on *Spartina alterniflora* and associated fungal growth, for which *L. irrorata* may act as a top-down control on plant growth. Yet, *L. irrorata* occupies other salt marsh plants, suggesting its habitat niche may be broader than previously reported. Here, we documented snail densities and size distributions in a Louisiana (USA) salt marsh composed of multiple marsh graminoids and report the results of behavioral choice experiments designed to test snail habitat preferences as a potential mechanism underlying their field distribution. We observed higher snail densities on *S. alterniflora* stalks (283.0 snails m^-2^) than other plant species, however, snails were highly abundant on *S. patens* (115.6 snails m^-2^), *Juncus roemerianus* (94.8 snails m^-2^), and *Distichlis spicata* (56.9 nails m^-2^) with densities comparable or higher on all species than reported on *S. alterniflora* in other studies along the U.S. Atlantic and Gulf coasts. Snails found on *S. alterniflora* and *J. roemerianus*, both plants with tall and rigid stalks, were also larger than snails found on other plant species. In species preference experiments, snails preferred *S. alterniflora* over *S. patens* and *D. spicata*, but no clear preferences were observed between *S. alterniflora* and *J. roemerianus*, nor between any combinations of *S. patens, D. spicata*, and *J. roemerianus*. Finally, we found that snails preferred senescing and dead *S. alterniflora* tissue over fresh *S. alterniflora*. Interpreting these results in tandem, this study suggests *L. irrorata* snails have consistent patterns of field distributions that match their habitat preferences, and future studies should test potential processes driving snail habitat selection, such as dietary habits and predator refugia (i.e., climbing sturdy stalks to avoid aquatic predators). Considering the abundance and trophic role of *L. irrorata* in coastal salt marshes, snail behavior may be a key modulator for salt marsh trophic networks.

## Introduction

Coastal salt marshes provide a variety of ecosystem services, including serving as a basal food source for a productive food web (McCann et al. 2017), providing structural refuge for juveniles of many commercially and recreationally important species (Peterson and Turner 1994, Able et al. 2015), protecting inland areas from high intensity storms (Costanza et al. 2008), and decreasing eutrophication via nutrient cycling and removal (Hopkinson and Giblin 2008). Marsh ecosystems can be stressful, owing to dynamic environmental conditions (e.g., temperature, salinity, inundation, dissolved oxygen, etc.) that can change rapidly. As a result, species assemblages include tolerant flora and fauna capable of withstanding extreme conditions. Marsh plants exhibit numerous adaptations for survival in these areas, including salt-excreting glands, resistance to flooding conditions, and broad thermal tolerances. Plant diversity within salt marshes can also be low, and this highlights the need for better understanding of the roles each plant species plays in supporting food webs. For example, omnivorous snails comprise highly abundant biomass pools and important trophic intermediates, facilitating the transfer of energy from basal plant production to higher trophic levels (Hamilton 1976, Silliman and Zieman 2001, McCann et al. 2017).

The salt marsh periwinkle snail (*Littoraria irrorata*) inhabits, and is often the dominant snail in, salt marshes of the Atlantic and Gulf coasts of the United States. This snail resides in emergent vegetation on the marsh platform where it displays distinct behavior of climbing plant stems at high tide to avoid predation by aquatic predators from below (Warren 1985, Carroll et al. 2018), as a mechanism of thermoregulation (Williams and Appel 1989, Henry et al. 1993), and to facilitate fungal invasion on plant leaves for subsequent consumption (Silliman and Newell 2003). *Littoraria irrorata* graze primarily on *Spartina alterniflora* compared to other plant species (e.g., Hendricks et al. 2011, Sieg et al. 2013), and especially graze senesced rather than live *S. alterniflora* leaves (Bärlocher and Newell 1994). *Littoraria irrorata* populations influence a variety of marsh ecosystem components including vegetation, microbial communities, organic matter and nutrient cycling, and marsh-estuarine food webs (Zengel et al. 2017). While some studies indicate *L. irrorata* exerts top-down control on plant aboveground biomass and productivity (Silliman and Zieman 2001), others have found no support for top-down control of marsh plant productivity (Kiehn and Morris 2009). Ecologists have most frequently associated study of *L. irrorata* with *S. alterniflora* that commonly define coastal salt marshes (e.g., Hamilton 1976, Silliman and Zieman 2001, Zengel et al. 2017, Rietl et al. 2018), and therefore the trophic role of *L. irrorata* has typically been considered linked (and largely limited) to the abundance and distribution of *S. alterniflora* (Silliman and Zieman 2001, McFarlin et al. 2015).

Coastal salt marshes are heterogeneous ecosystems containing mosaics of plant species arranged in patches that reflect variation in local conditions (e.g., elevation/inundation frequency, salinity, soil properties, etc.) and interspecific competition (Pennings et al. 2005). In addition to the dominant *S. alterniflora*, coastal marshes along the United States Gulf Coast also contain patches of the macrophytes *S. patens, Juncus roemerianus*, and *Distichlis spicata*. Salt marsh plants in this region exhibit different responses to hydrologic alterations including flooding frequency and salinity stress (Jones et al. 2016). As a result, future changes in climate, inundation, and salinity regimes are predicted to change plant community structure in coastal salt marshes. Given this, developing a more comprehensive understanding of plant-animal interactions involving abundant marsh plants is critical information for predicting future food webs and trophic structure.

High densities of *L. irrorata* snails have been reported in salt marshes along the Gulf and Atlantic coasts of the United States (Rietl et al. 2018). However, few studies have quantified the densities or lengths of snails on different types of salt marsh vegetation (but see Hughes 2012, Faillon et al 2020). Further, little information is available on snail preferences for different vegetation types that vary in relative abundances across the marsh landscape. Here, we used empirical and experimental investigations to determine the abundance and habitat preferences of *L. irrorata* snails across the marsh landscape. We provide field-based estimates of snail density and size distributions across multiple plant species in a Louisiana salt marsh. In addition, we performed controlled laboratory experiments to test snail preferences for: 1) various species of marsh plants, 2) stages of *S. alterniflora* senescence, and 3) plant tissue against structural controls. We predicted snail field distributions would be reflected in experimental choice tests and that snails would prefer senesced over live plants and live plants over wooden structural controls. The overarching goal of this research was to gain insight into snail distributions and preference patterns to develop a better understanding of the marsh ecosystem and food web.

## Materials & Methods

### Snail Densities and Size Distributions

We quantified snail densities and lengths at a well-studied marsh site (Able et al. 2015, Marton et al. 2015, Bernhard et al. 2019, Rietl et al. 2018, Keppeler et al. 2021) along the northwestern shore of Bay Batiste (29.4759°N, 89.8543°W) on seven (snail length) to nine (snail density) dates between May 2016 and January 2018 (Table S1). On each date, we collected snails in at least three replicate 0.25 × 0.25 m quadrats from within each of four monoculture vegetation types (*S. alterniflora, S. patens, D. spicata*, and *J. roemerianus*). All snails were kept on ice during transport, and stored at 4°C to await processing, which was completed within 48 h of collection. In the laboratory, we rinsed, cleaned, and counted all snails before measuring shell length (mm) using digital calipers, which had accuracy and precision of 0.01 mm.

### Habitat Preference Experiments

We experimentally determined snail habitat preferences following methods established in previous habitat choice studies (Martin 2017, Martin et al. 2020). All trials were performed in 20-liter (15cm x 30cm x 20cm) arenas, each containing 100 mL of 9 psu seawater (Instant Ocean^®^, Instant Ocean Spectrum Brands), a typical salinity for Gulf of Mexico salt marshes. We obtained snails and plants from marshes near the Louisiana Universities Marine Consortium (LUMCON)’s DeFelice Marine Center in Cocodrie, Louisiana (USA) (29.2580°N, 90.6629°W), a representative coastline with extensive salt marsh (e.g., Hill and Roberts 2017). Snails were collected from stands of the four studied species of vegetation and stored in the same cooler for transport so that the source vegetation was randomized. We performed three experiments to test snail preference patterns for: 1) marsh plant species, 2) *S. alterniflora* state of senescence, and 3) plant matter as opposed to structural controls. In experiment 1, we tested snail preference for plant species by offering a choice between each of the following combination of plants (n=10 for each combination): *S. alterniflora* vs. *S. patens, S. alterniflora* vs. *D. spicata, S. alterniflora* vs. *J. roemerianus, S. patens* vs. *D. spicata, S. patens* vs. *J. roemerianus*, and *D. spicata* vs. J. *roemerianus*. In experiment 2, we tested snail preference for different *S. alterniflora* states of senescence, using all combinations of green (live), yellow (partially senesced), and brown (dead) *S. alterniflora*, as described by Graça et al. 2000 (n=10 for each combination). Finally, experiment 3 was conducted to determine whether snails preferred any of the four plant species (*S. alterniflora, S. patens, D. spicata*, and *J. roemerianus*) to a structural control, which offers a rigid structure to climb and escape aquatic predators, but no viable food source (n=5 for each combination).

We cut all plants used in experiments to 15-cm segments (the height of the arena) using only tissue from between the first leaf and final leaf of a stem for *S. alterniflora, S. patens*, and *D. spicata* samples, and using tissue at least 10 cm above the exposed base of a blade and at least 10 cm below the tip of a blade for *J. roemerianus*. We rinsed and mounted plant stalks in polystyrene foam inserts and placed them at each end of the arena. All stalks were standardized to contain equal volume (90 cm^3^) of each species used in trials (4-15 stalks used per trial). In experiment 3, we used six dowels of approximately equal diameter (1 cm) and height (15 cm) as paired plant stalks to offer structural refuge but no viable food source paired with one of the four plant species.

We randomly selected six snails (20-25mm shell length) for use in each trial after starving snails for 48 h. This density is within the natural range of snail densities we observed in this study and reported in Reitl et al. (2018). We placed snails in the middle of arenas, and a camera mounted 30 cm above each arena captured a photograph of the arena interior at five-minute intervals for 12 hours. Due to this short trial duration, controls to correct for autogenic or allogenic changes to plant tissues were not necessary, as the amount of decomposition of plant tissue in 12 hours was minimal (Roa 1992). We covered arenas with Saran^®^ wrap to prevent snail escape while maintaining visibility from above for time-lapse photography. Each trial included six hours of simulated daylight (four white fluorescent lights at 25°C) and six hours of simulated night (four low-wattage violet fluorescent lights at 20° C) to account for diurnal differences in snail behavior (Graça et al. 2000, Iacarella and Helmuth 2011). To capture snail behavior during night conditions, we marked snail shells with odorless neon fluorescent paint prior to placement in arenas (Fig. S1). We began half of trials as simulated day, and the other half as simulated night, and the order did not affect snail habitat preference (Kruskal-Wallis H_2, 356_ = 178.4, *P* = 0.295).

We processed snail preferences using time-lapse photography, which we recorded at five-minute intervals to quantify preference patterns (Video S1). We defined a preference for a habitat by a snail as any part of the snail contacting a plant or wooden dowel, or if a snail was attached to an arena wall within 0.5 cm of a plant’s canopy or a wooden dowel. We calculated the amount of time each of the six snails per trial exhibited a preference for one available habitat versus the opposite available habitat, and then averaged across all time points for a trial (144 time points per trial) to derive the proportion of time exhibiting preference. For example, if six snails on average spent 40% of a trial within the *S. alterniflora* habitat, and 5% of a trial within the *D. Spicata* habitat, this would suggest a preference for *S. alterniflora*.

### Statistical Analyses

In the field survey, a preliminary analysis indicated that snail density and length varied little across sampling dates (one-way ANOVA: *p* > 0.05). As a result, we pooled dates and conducted analyses using only plant species as the predictor variable. Time-pooled snail densities and lengths were both normally distributed. We conducted separate one-way analysis of variance (ANOVA) for response variables of snail density (snails m^-2^) and length (mm).

For each choice comparison in experiments, we evaluated differences in preference between habitats by conducting a matched pairs t-test of the difference in the proportion time spent within each provided habitat. We arcsine transformed all proportion data derived from habitat preference experiments in order to stabilize variance and reduce the dependency of variance upon the mean, to uphold assumptions of normality for paired t-tests (Sokal and Rohlf 1995). We conducted all statistical analyses in R 3.5.0 (R Core Team 2018) and with use of the *tidyverse* package (Wickham et al., 2019) and considered all results significant at *p* < 0.05.

## Results

### Snail Densities and Size Distributions

We found *L. irrorata* in plots within each of the four salt marsh vegetation types (*S. alterniflora, S. patens, J. roemerianus*, and *D. spicata*) in Bay Batiste, Louisiana on all nine sampling dates between May 2016 and January 2018. Snail density did not differ with time within any of the vegetation types (Table S1). Across all sampling dates, snail density was significantly (*p* < 0.05) higher on *S. alterniflora* (mean ± SE = 283.0 ± 29.4 snails m^-2^) than the other three vegetation types (Fig. 1a), and densities on *S. patens* (115.6 ± 25.8 snails m^-2^) and *J. roemerianus* (94.8 ± 26.2 snails m^-2^) were 2.4 – 3.0 times lower than *S. alterniflora* and 1.7 – 2.0 times higher than densities on *D. spicata* (56.9 ± 10.0 snails m^-2^). Individual snail shell lengths were significantly larger (*p* < 0.05) for snails collected in *J. roemerianus* (22.02 ± 0.14 mm) and *S. alterniflora* (21.84 ± 0.08 mm) than *D. spicata* (20.99 ± 0.22 mm) and *S. patens* (20.98 ± 0.14 mm) (Fig. 1b).

**Figure 1.**
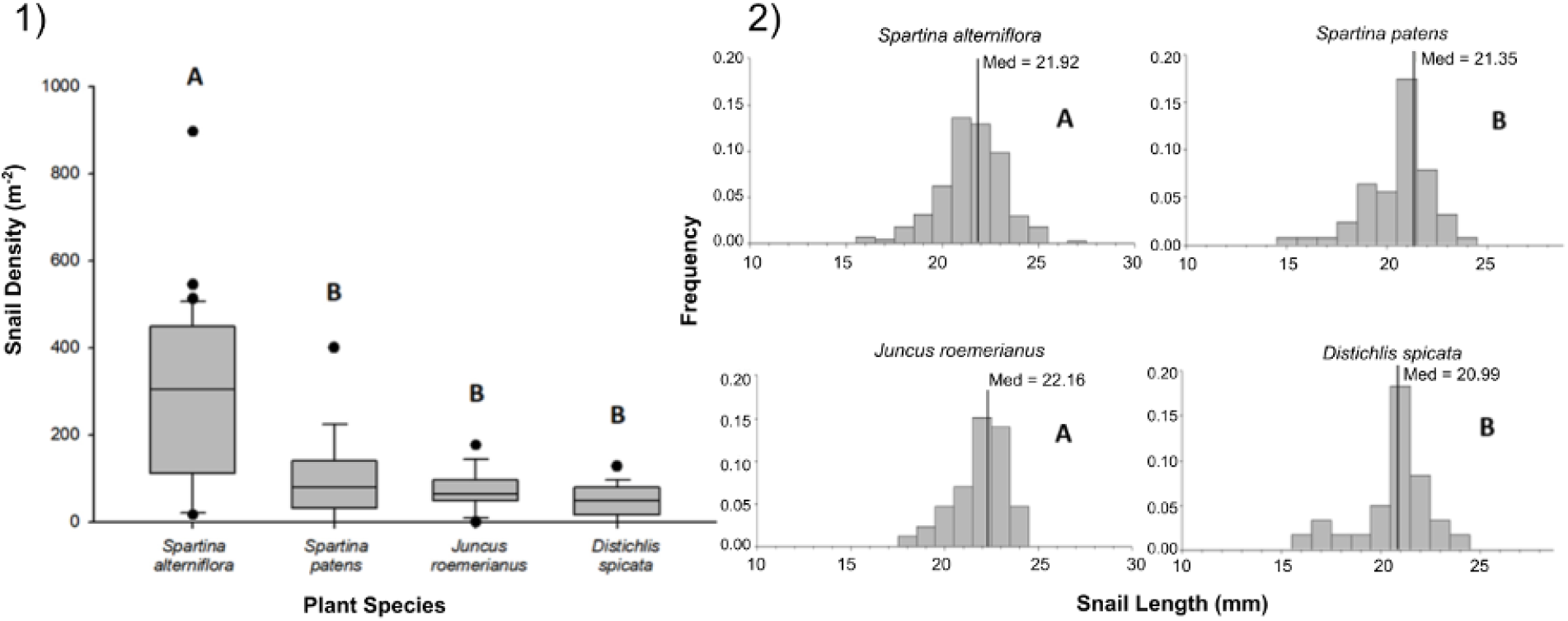
Density and size of periwinkle snails on marsh plant species. 1) Density of snails in 0.25m x 0.25m plots of *Spartina alterniflora, Spartina patens, Juncus roemerianus*, and *Distichlis spicata* in a salt marsh in Bay Batiste, LA. Values derived from 3 -5 plots per vegetation type on nine dates between May 2016 and January 2018. Boxplots show median, 25th and 75th percentiles (lower and upper hinges, respectively), and whiskers extend to largest and smallest values, unless values are greater or less than 1.5 * IQR (outlying points plotted individually). 2) Length (mm) distributions for snails collected on each of the four vegetation species on the same nine dates. Vertical lines represent the median values for each vegetation type. In both panels, different letters indicate statistically significant (p < 0.05) differences among vegetation types. Snail densities were higher in *S. alterniflora* stands than in other vegetation, and snails were longer in *S. alterniflora* and *J. roemerianus* stands than *S. patens* and *D. spicata* stands.

### Experiment 1: Species Preference

*Littoraria irrorata* snails used in this experiment exhibited clear and significant preferences between marsh plant species (Fig. 2, Table 1). Snails occupied *S. alterniflora* more often than two of the three other common marsh plant species: on average, we found snails on *S. alterniflora* 13.6 and 4.3 times more often than on *S. patens* and *D. spicata*, respectively.

**Table 1.**
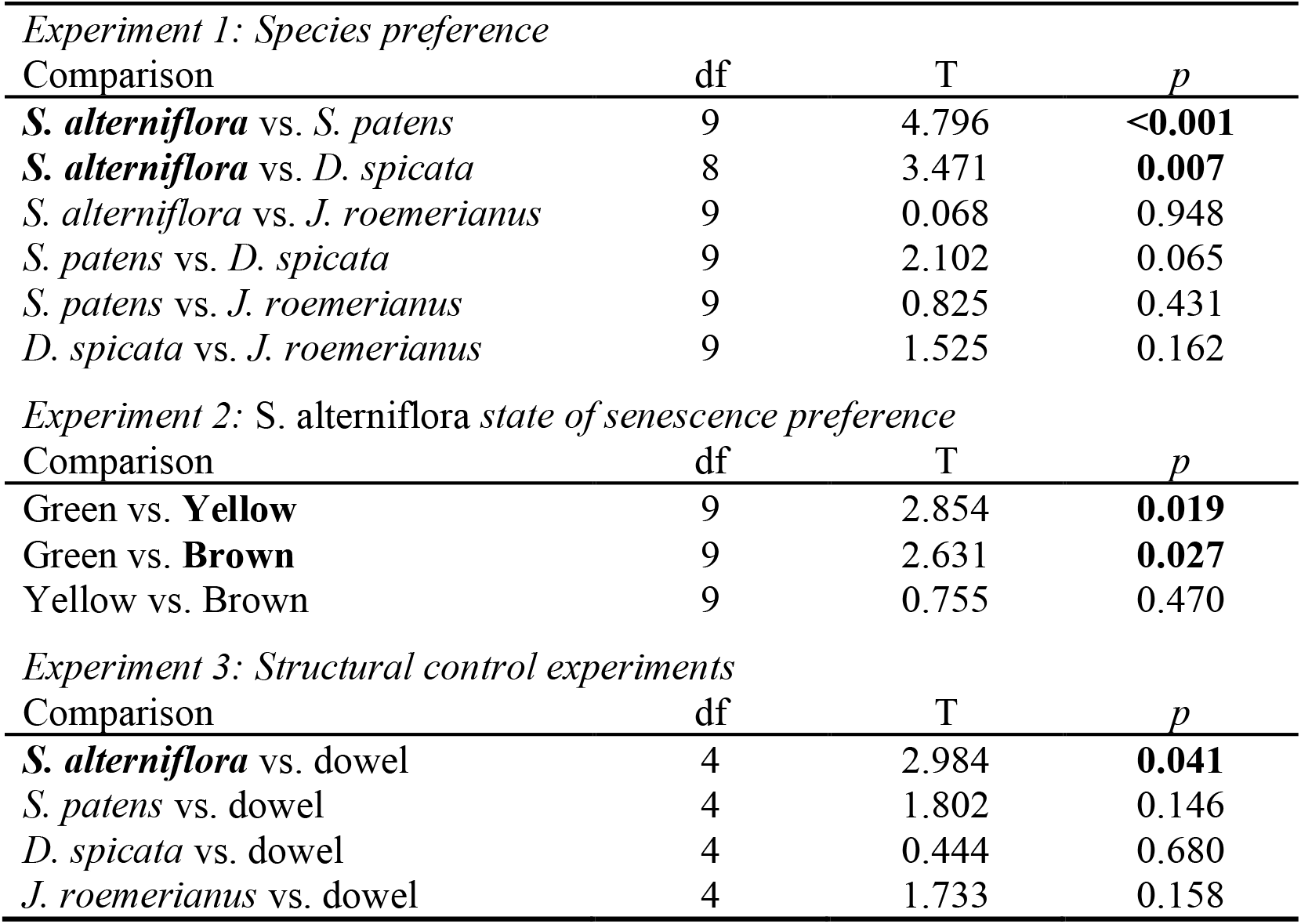
Statistical results from matched pairs t-tests for species, *S. alterniflora* state of senescence, and structural control preference experiments. P-values < 0.05, as well as the option of the pair for which a significant preference was demonstrated, are denoted in bold.

| *Experiment 1: Species preference* |  |  |  |
| --- | --- | --- | --- |
| Comparison | df | T | *p* |
| *S. alterniflora* vs. *S. patens* | 9 | 4.796 | **<0.001** |
| *S. alterniflora* vs. *D. spicata* | 8 | 3.471 | **0.007** |
| *S. alterniflora* vs. *J. roemerianus* | 9 | 0.068 | 0.948 |
| *S. patens* vs. *D. spicata* | 9 | 2.102 | 0.065 |
| *S. patens* vs. *J. roemerianus* | 9 | 0.825 | 0.431 |
| *D. spicata* vs. *J. roemerianus* | 9 | 1.525 | 0.162 |
| *Experiment 2: S. alterniflora state of senescence preference* |  |  |  |
| Comparison | df | T | *p* |
| Green vs. **Yellow** | 9 | 2.854 | **0.019** |
| Green vs. **Brown** | 9 | 2.631 | **0.027** |
| Yellow vs. Brown | 9 | 0.755 | 0.470 |
| *Experiment 3: Structural control experiments* |  |  |  |
| Comparison | df | T | *p* |
| *S. alterniflora* vs. dowel | 4 | 2.984 | **0.041** |
| *S. patens* vs. dowel | 4 | 1.802 | 0.146 |
| *D. spicata* vs. dowel | 4 | 0.444 | 0.680 |
| *J. roemerianus* vs. dowel | 4 | 1.733 | 0.158 |

**Figure 2.**
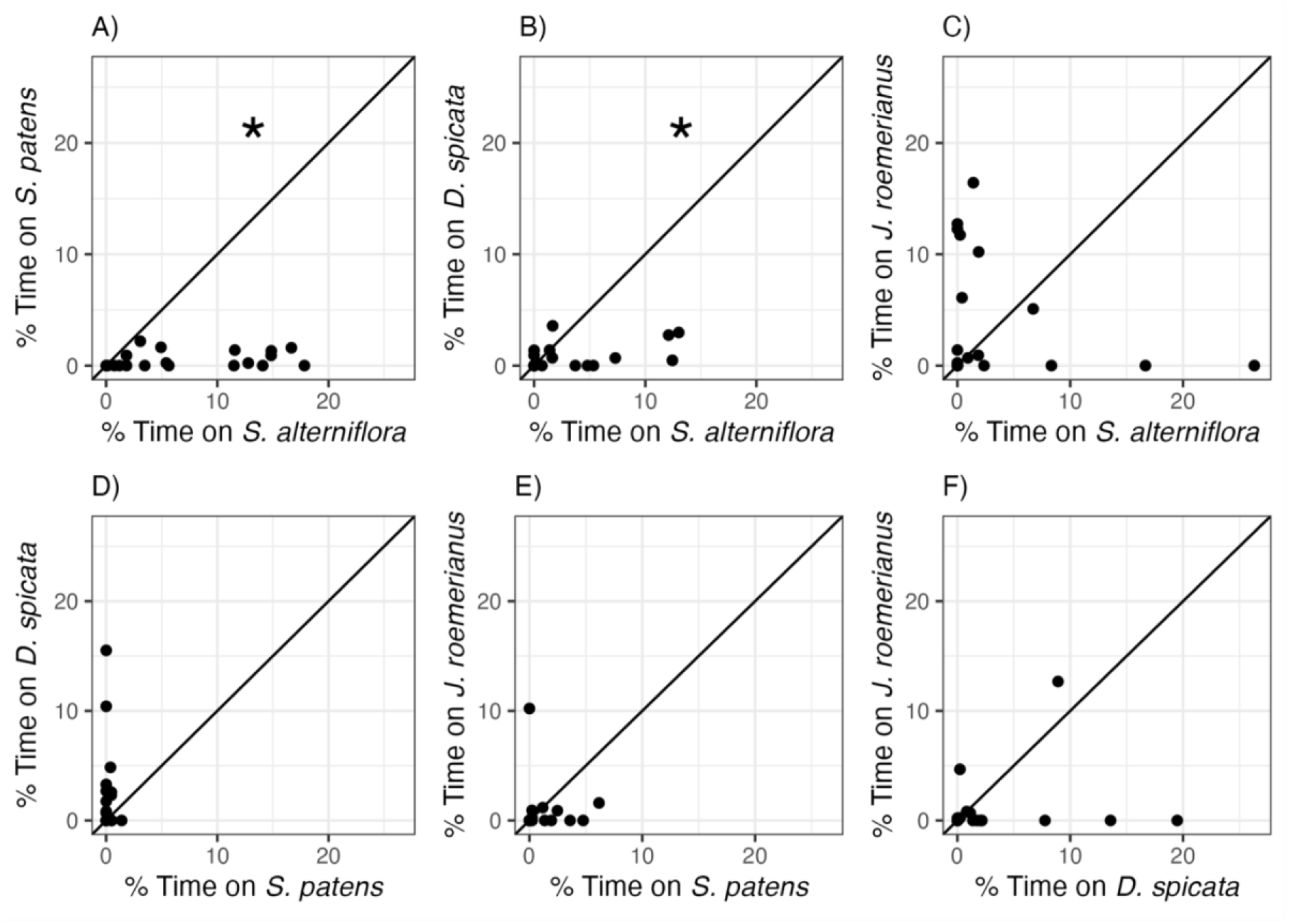
Snail preferences for different marsh plant species. Percent of time snails spent on A) *S. patens* vs. *S. alterniflora*, B) *D. spicata* vs. *S. alterniflora*, C) *J. roemerianus* vs. *S. alterniflora*, D) *D. spicata* vs. *S. patens*, E) *J. roemerianus* vs. *S. patens*, and F) *J. roemerianus* vs. *D. spicata* in choice experiments. Each point represents mean time spent within each habitat type by six snails within each choice trial. Line represents the 1:1 line which indicates no preference between the two choices. Statistically significant preferences (p < 0.05) are indicated with *.

However, there was no significant difference in time spent on *S. alterniflora* (3.35 ± 6.79 %) and time spent on *J. roemerianus* (3.90 ± 5.56 %). We also found snails 12.2 times more often, on average, on *D. spicata* (2.25 ± 4.01 %) than on *S. patens* (0.183 ± 0.348 %; Fig. 2d), but this difference was not significantly different (*p* = 0.060, Table 1). Snails did not show a significant preference for *J. roemerianus* compared to either *S. patens* or *D. spicata* (Fig. 2e,f).

### Experiment 2: S. alterniflora State of Senescence Preference

When snails were given a choice between *S. alterniflora* stems at different stages of senescence, snails significantly preferred partially senesced or dead *S. alterniflora* stems over live stems (Fig. 3., Table 1). Snails were observed on yellow (partially senesced) and brown (dead) stems 7.0 and 3.2 times more frequently, on average, than on green (live) stems (Fig 3a,b). There was no consistent or significant difference in the proportion of time spent by snails on yellow (partially senesced) (13.7 ± 16.1%) compared to brown (dead) (9.8 ± 9.0%) stems (Fig. 3c.)

**Figure 3.**
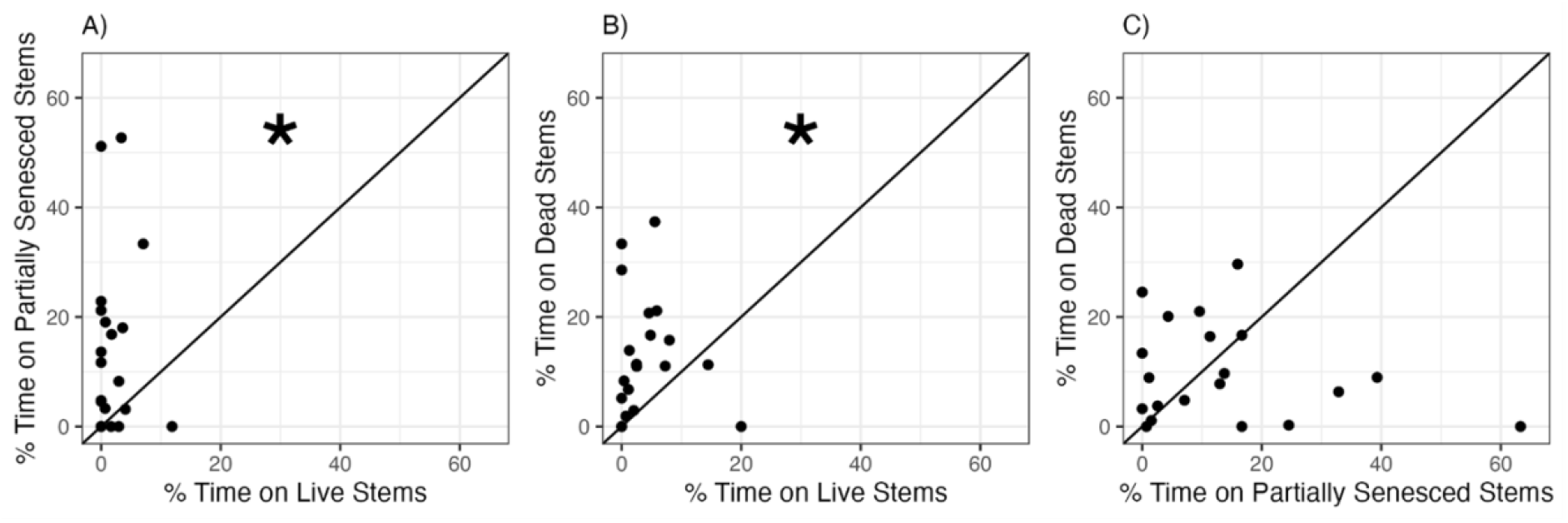
Snail preferences for *Spartina alterniflora* at different stages of senescence. Percent of time snails spent on A) yellow (partially senesced) vs. green (live), B) brown (dead) vs. green (live), or C) brown (dead) vs. yellow (partially senesced) *Spartina alterniflora* stems in choice experiments. Each points represents mean time spent within each habitat type by six snails within each choice trial. Line represents the 1:1 line which indicates no preference between the two choices. Statistically significant preferences (p < 0.05) are indicated by an asterisk

### Experiment 3: Structural Control

When snails were given a choice between plant stems and a structural control (wooden dowels), snails exhibited a variable preference response depending on which plant species was offered. Snails were 60 times more likely to be found on *S. alterniflora* (13.8 ± 19.8%) than on structural controls (0.2 ± 0.5%) (Table 1). In contrast, snails did not show a consistent or significant preference for any of the other three plant species over the structural controls (Table 1).

## Discussion

As one of the most common organisms in the salt marshes of North America and a key trophic link in these ecosystems, the marsh periwinkle snail *L. irrorata* plays a significant role in salt marsh nutrient and energy flow (Silliman and Bertness, 2002). Although *L. irrorata* are thought to be dietary specialists for the smooth cordgrass *Spartina alterniflora* and fungal growth on this plant (Silliman & Zieman, 2001), they are also found on several other plant species (Hughes 2012, Failon 2020), and their preferences between host plants remain unclear. Here, we combined field observations of snail densities among four common salt marsh graminoids– *S. alterniflora, S. patens, D. spicata*, and *J. roemerianus–* on nine sampling dates over a 20-month period with experiments of habitat choice among the same four species, to explore links in snail behavior and distributions.

Across almost two years of field surveys in Bay Batiste along the Louisiana Gulf Coast, snails were highly abundant on all four plant species, with densities comparable or higher on all species than reported on *S. alterniflora* in other studies along Atlantic and Gulf Coasts (McFarlin et al. 2015, Rietl et al. 2018). Furthermore, high densities were consistent across seasons, suggesting the persistence of snails within stands. This reflects spatial persistence of snails in Florida (Hamilton 1978) and the Gulf coast in Texas (Vaughn and Fisher 1992). Several prior studies of *L. irrorata* distributions reported only high snail densities *S. alterniflora* stands (e.g. Watson and Norton 1985, Silliman and Zieman 2001). However, our findings more closely reflect those of Hughes (2012), who found that *L. irrorata* densities were highest in mixed stands of *S. alterniflora* and *J. roemerianus*, and Failon et al. (2020), who found comparable snail densities in adjacent stands of *S. alterniflora* and *S. cyrosuroides*. High *L. irrorata* densities in plant stands composed of species other than *S. alterniflora* suggests a broader habitat niche for the snail than previously assumed, motivating experimental examination of habitat preferences across plant species. Furthermore, these high densities may amplify *L. irrorata*’s importance in salt marsh trophic networks.

In species preference experimentation, *L. irrorata* demonstrated a significant preference for *S. alterniflora* over *S. patens* and *D. spicata*, but no clear preferences between *S. alterniflora* and *J. roemerianus*, nor between any combinations of *S. patens, D. spicata*, and *J. roemerianus*. Snail preference for *S. alterniflora* over other species of plant was expected, as *L. irrorata* was most abundant in *S. alterniflora* patches in the field. There may be several mechanisms underlying this preference, however. One possibility, as suggested in prior studies, is that *S. alterniflora* is a known food source for *L. irrorata*. Yet *L. irrorata* also derives nutrition from epiphytic microalgae (which can grow on the stalks of many plant species), and benthic algae and detritus in the marsh soils (Alexander 1979, Watson and Norton 1985). Furthermore when food sources are plentiful, such as is often the case for *L. irrorata*, snail distributions may be determined by other criteria, such as predator avoidance (Zaret and Suffern 1976, Loose and Dawidowicz 1994). *Callinectes sapidus*, the primary predator of *L. irrorata*, is a threat from below that patrols the marsh tidal zone, but cannot access snails higher in the marsh canopy (Hughes 2012). Thickness and rigidity of stalks may therefore play a considerable role in habitat selection when snail densities are high; a frail plant may bend or collapse under the weight of many snails. Of the four plant species studied here, *S. alterniflora* has the widest average thickness (Hester, Mendelssohn & McKee, 2001), yet *J. roemerianus* has the most rigid stems (Eleuterius, 1976). Such attributes may explain the lack of a significant difference in time spent on *S. alterniflora* and *J. roemerianus*. While there is no evidence that *J. roemerianus* tissue serves as a food source for *L. irrorata, L. irrorata* may persist in *J. roemerianus* stands by grazing on epiphytic microalgae, benthic algae, and detritus. Conversely, both *S. patens* and *D. spicata* have thin stalks that, on several occasions observed during choice experiments, collapsed under the weight of multiple snails. Refuge-seeking behavior on sturdy stalks of *S. alterniflora* and *J. roemerianus*, particularly for larger snails, also would explain empirical snail observations on the four plant species: snail densities were not only higher, but snail shell lengths were also longer, in stands of *S. alterniflora* and *J. roemerianus*. Along with stem thickness and rigidity, stem height may influence host selection and predator avoidance by snails (Hughes 2012) with *S. alterniflora* and *J. roemerianus* typically growing taller than *S. patens* and *D. spicata* in these habitats. Plant stem height was standardized within and across species in these experiments, but the combination of stem thickness, rigidity, and height may influence snail refuge-seeking behaviors in the field.

Widespread removal of snail predators may dramatically impact snail habitat preferences and grazing behaviors if these preferences are motivated by predator avoidance. *Callinectes sapidus* populations on the eastern seaboard of the USA declined significantly in the 20^th^ century up to present day (Abbe and Stagg 1996, Cole 1998, Kahn et al. 1998, Lipcius and Stockhausen et al. 2002, Lycett et al. 2020), and a number of physical and anthropogenic factors influence crab distributions (Jivoff et al. 2017). Changes in crab population structure likely affect snail distributions between marsh graminoid taxa and across vertical (ground-to-canopy) gradients.

However, *L. irrorata* may still express predator avoidance behaviors even if no predators are present (Hughes 2012), which suggests that *L. irrorata* may select for rigid plant stalks even if predator abundance is low. Future work should be aimed at determining the relative roles of snail foraging and predation in governing snail occupancy patterns.

We explored *L. irrorata* preferences between live, partially senesced, and fully senesced *S. alterniflora* stalks to complement choice experiments of live-only plant stalks from four species. Here, snails spent more time on partially senesced and standing dead *S. alterniflora* than on live green *S. alterniflora*, but snails did not exhibit a preference between partially senesced and standing dead plant tissue. These findings are consistent with our predictions and prior evidence on snail litter preferences (Bärlocher and Newell 1994). Experimental habitat preference is thought to be a good indicator of associated grazing behaviors in the field (Leighton 1966, Keesey et al. 2015), and in 23 out of 65 trials involving any form of *S. alterniflora* tissue there was evidence of grazing (long radulations on tissue) after just twelve hours of trial (Fig. S2). While snails in these experiments may have selected host plants due to factors beyond grazing quality, our experimental results of snail habitat preferences here are consistent with our predictions, and prior work that showed *L. irrorata* graze more upon fully senesced *S. alterniflora* tissue than live *S. alterniflora* tissue (Bärlocher and Newell 1994). Previous analyses of snail stomach materials found that less than 2% (Silliman and Zieman 2001) and 3% (Alexander 1979) of snail gut content was live green plant material. Grazing experiments conducted by Bärlocher and Newell (1994) suggested a preference for standing dead leaves, in both recently collected and powdered form, over respective forms of “yellow-green” leaves (defined as 25-30% green tissue remaining). Senescing *S. alterniflora* tissue has higher lipid content and concentrations of desired fungal epiphytes than green tissue, and soft, decaying tissue is easily digested compared to live tissue (Bärlocher and Newell 1994, Silliman and Zieman 2001).

Synthesizing results from our field studies and choice experiments in the context of prior work, it remains possible that bottom-up (food availability and quality), top-down (predator avoidance), or both sets of factors may drive snail decision-making and distributions. We speculate that snails may therefore face a series of hierarchical decisions in selecting a plant host: if food availability is low, preference may be exhibited for *S. alterniflora*, especially decaying or dead tissue. When food availability is high, snails may seek out tall or rigid stems (e.g. *J. roemerianus*) that may best serve as refuge from aquatic predators (Hughes 2012). High snail densities on all four plant species, combined with selection for *J. roemerianus* at comparable rates as *S. alterniflora*, also lends evidence to a broader habitat niche for the snail than previously suspected. Acting as a habitat generalist, rather than interacting only with a single plant species, may indicate wider recruitment across the marsh platform. Given that adult *L. irrorata* do not disperse far beyond where they have passively settled in their planktonic larval form (Hamilton 1978, Vaughn and Fisher 1992), tolerance of multiple plant hosts may enable snail colonization of mixed-vegetation habitats. Furthermore, a broader habitat niche may also offer snails greater resiliency in the face of disturbance, such as salinity changes or climate change-induced sea level rise, which can alter the composition of plant communities (Morris et al., 2002). Persistence of snails in the face of disturbance is important given their central connectedness to the rest of the marsh food web. Yet given the strong preferences of the snail for decaying *S. alterniflora* tissue, presumably for high forage quality, broader habitat preferences may also create an ecological trap. If snails select for, and remain in, monotypic stands of *J. roemerianus* to avoid predation, they may experience lower food availability or nutritional content, as their grazing is limited to microalgae and detritus rather than plant tissue. Snail decision-making in light of multiple possible hosts may therefore both provide plasticity and vulnerability, depending on the heterogeneity of available plant hosts.

## Conclusions

Animal behavior is an important process that influences habitat preferences and can determine the distribution of organisms in space and time. Here, we quantified natural densities of the marsh periwinkle *L. irrorata* on four common marsh graminoids, and explored the mechanisms driving such observations via habitat choice experiments. Our findings suggest that the habitat niche of *L. irrorata* may be broader than single-species specialization for *S. alterniflora*. Broad habitat tolerances may provide the snail greater resilience to the stressful, dynamic conditions of salt marshes, if both food sources and adequate predator refugia are available. Given the high abundance of this marsh omnivore, its behaviors likely have important implications for salt marsh nutrient and energy flow. With a habitat preference for decaying plants over live tissue, *L. irrorata* may control plant productivity only during peak growth when senesced tissue abundance is low. We suggest a key modulating role of snail behavior for the salt marsh trophic network, drawing upon our combination of empirical observations with preference experiments.

## Acknowledgements

We thank the members of the Roberts lab, especially Ron Scheuermann, Madelyn Sorrentino, Nicole Farley, Jacqueline Levy, Ariella Chelsky, and Anthony Rietl, for assistance in preliminary field lab work, and the LUMCON community for a productive research atmosphere during an REU experience. We thank the Department of Biological Sciences at Dartmouth College, especially Drs. Matt Ayres, Celia Chen, and Hannah ter Hofstede for guidance and constructive criticism.

---

**Table S1.** *Littoraria irrorata* field densities across time. Mean + standard error (SE) of *Littoraria irrorata* densities (snails m^-2^) in 0.25m x 0.25m single-species plots (n = 3 -5) of *Spartina alterniflora, Spartina patens, Juncus roemerianus*, or *Distichlis spicata* on each of nine sampling dates in a salt marsh in Bay Batiste, LA.

| Date | *S. alterniflora* |  | *S. patens* |  | *J. roemerianus* |  | *D. spicata* |  |
| --- | --- | --- | --- | --- | --- | --- | --- | --- |
|  | Mean | SE | Mean | SE | Mean | SE | Mean | SE |
| 5/16/2016 | 345.6 | 68.0 | 96.0 | 66.6 | 112.0 | 24.4 | 64.0 | 16.0 |
| 6/15/2016 | 341.3 | 85.8 | 272.0 | 84.7 | 288.0 | 9.2 | 117.3 | 32.4 |
| 8/16/2016 | 293.3 | 129.4 | 117.3 | 64.9 | 101.3 | 23.2 | 32.0 | 9.2 |
| 10/11/2016 | 269.3 | 91.2 | 90.7 | 50.9 | 26.7 | 14.1 | 48.0 | 16.0 |
| 2/1/2017 | 406.4 | 147.7 | 64.0 | 16.0 | 32.0 | 24.4 | 58.7 | 29.7 |
| 5/9/2017 | 176.0 | 78.9 | 202.7 | 101.3 | 85.3 | 19.2 | 53.3 | 14.1 |
| 9/7/2017 | 117.3 | 32.4 | 85.3 | 10.7 | 53.3 | 23.2 | 80.0 | 33.3 |
| 10/26/2017 | 293.3 | 75.2 | 10.7 | 10.7 | 96.0 | 40.2 | 48.0 | 24.4 |
| 1/25/2018 | 304.0 | 99.9 | 101.3 | 29.7 | 58.7 | 10.7 | 10.7 | 10.7 |
| **Mean** | **283.0** | **29.4** | **115.6** | **25.8** | **94.8** | **26.2** | **56.9** | **10.0** |

**Figure S1.**
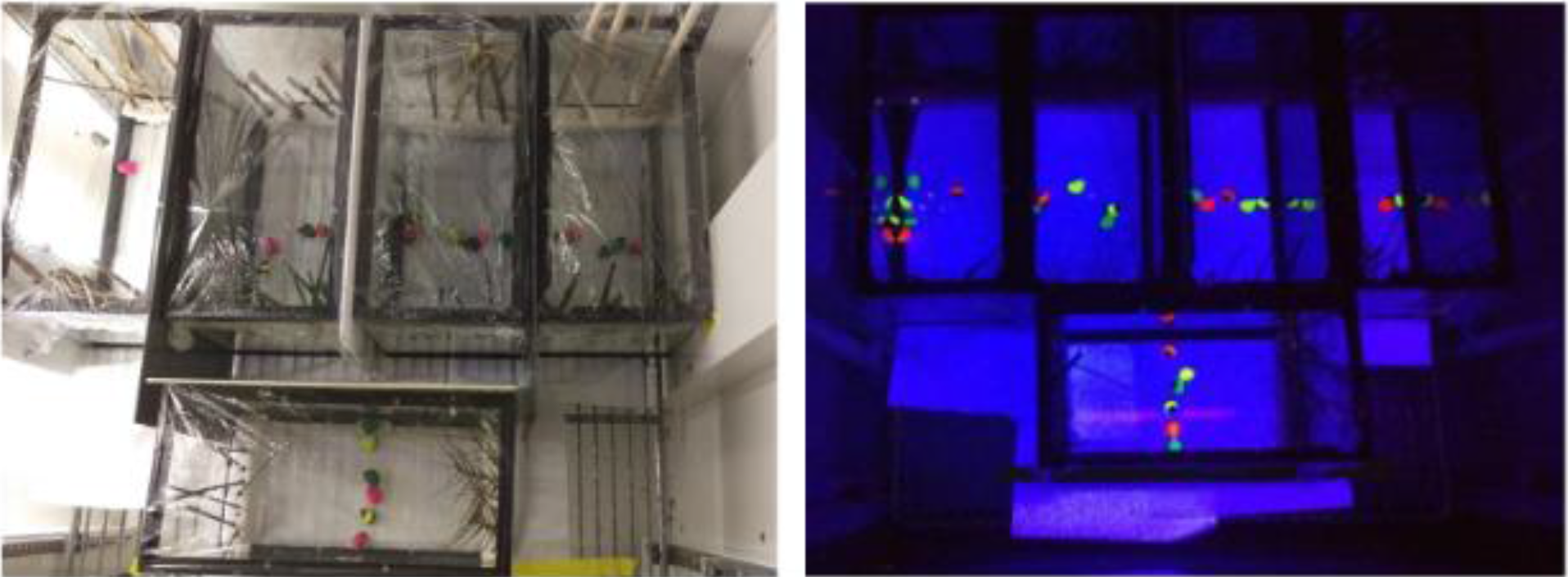
Aquaria design for habitat preference analysis, displayed during simulated daytime (left) and nighttime (right) conditions. Sets of *S. alterniflora*, S. *patens, D spicata, J. roemerianus* and wooden dowels were mounted on opposite sides using Styrofoam inserts, and six snails were placed in the center of each arena. Cameras were mounted above to record snail movement.

**Figure S2.**
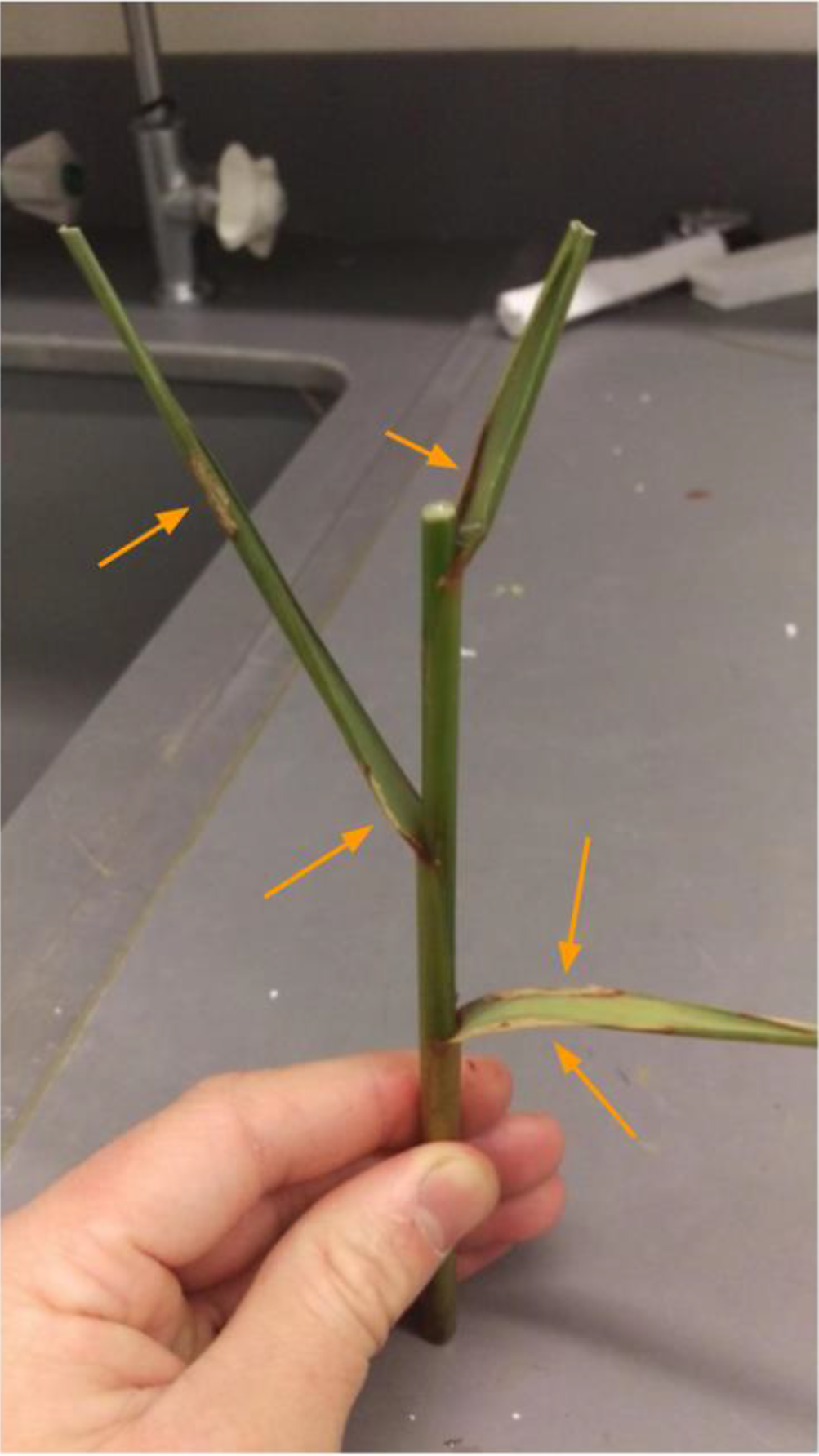
Evidence of grazing upon *S. alterniflora*. Radulations were concentrated on leaves of *S. alterniflora* plants, and were observed on many of the plant segments used in experimentation.

## Notes

### Competing Interest Statement

The authors have declared no competing interest.

